# The anticancer activity and immune modulating properties of combinatorial ensemble complementary (antisense) microRNA (fRNA) in combination with the immunomodulator - glycoproteid lectin from *B. subtilis* B-7025

**DOI:** 10.1101/151829

**Authors:** Artur Martynov, Gennady Didenko, Boris Farber, Sophya Farber, Olena Cruts

## Abstract

**Background:** Many adenocarcinomas have the ability to capture from an extracellular matrix the oligonucleotides and nanoparticles by pinocytosis, when the non-cancerous cells are not capable to capture the oligonucleotides and small liposomes. This provides selective accumulation of proposed protected oligonucleotides (fRNA) in cancer cells and also provides the absence toxicity in the fRNA.

**Materials and methods:** For the immunotherapy we used immunotropic 70 kDa lectin *B. subtilis B-7025*. *In vivo* experiments were carried out in C57BL line mice in Lewis lung carcinoma. The cytotoxic activity, lymphocytes, macrophages was determined *in vitro* using the MTT assay.

**Results and discussion:** Animal survival rate in groups receiving either the fRNA or Vaccine was 70 and 40%, respectively. Combined use fRNA and Vaccine has the advantage compared with the use these drugs in monotherapy, as the anticancer efficacy of the scheme is much higher, which is manifested in the primary tumor node and metastasis inhibition.

## Introduction

One of the new trends in the cancer treatment is the development of the gene cancer therapy [1]. Genes inactivation based on the antisense polynucleotides (AP) is one of the most perspective gene therapy directions [2]. The main problems to the implementation for anticancer gene therapy is the destruction of synthetic complementary nucleotides hydrolysis by blood nucleases, the limited penetration capability into cells, sensitivity to cell genome repair systems [3].

There are different approaches to the antisense oligonucleotides design, but the principle of their interaction with the targets remains the same: the hydrogen bonds formation between complementary nucleotides and increasing the degree of stability to nucleases. In some cases, developers protected the 5′-oligonucleotide from nucleases, in others modified 3′- [4]. Researchers from the US have replaced the deoxyribosyl residues to morpholine fragments aiming to create RNA/DNA fragments, resistant to nucleases [5]. At the same time, the gene inactivation principle through their interaction with the complementary antisense-nucleotides (hydrogen bonding) remained unchanged. Complementary microRNA with selectively blocking the synthesis of certain proteins in the cell are also known.

Many adenocarcinomas have the ability to capture from an extracellular matrix the oligonucleotides and nanoparticles by pinocytosis [6], both as liposomes [7] and the metal particulate forms [8,9,10]. However, non-cancerous cells are not capable to capture small oligonucleotides and liposomes [11]. This supports selective accumulation of the proposed protected oligonucleotides (fRNA-fragmented modified alanine tRNA) in cancer cells and also provides the absence toxicity in the fRNA preparation. For synthesis of fRNA we used 8 fragments of alanine tRNA, which gained the complementarity and endonuclease stability through succinylation of their structure on exocyclic amino groups of adenine and guanine as well as alcohol-free ribose group.

The selective accumulation of fRNA in the cancerous cell results to its hybridization with complementary targets in matrix alanine tRNA of cancerous cells and leads to a gradual stop of protein synthesis due to its blockade in the phase of alanine incorporating into the polypeptide chains. The mechanism of several oligonucleotides or oligopeptides interaction with a large target on insulin model as an example is presented in the following reference [12]. The fRNA effect is based on apoptosis induction through a stop of protein synthesis. Actually, fRNA has no effect on healthy cells, blocking the synthesis of all cellular proteins and excluding tumor adaptation to the anticancer therapy and selection of resistant cancer cells.

Another perspective direction in the development of anticancer drugs is selective cellular immunity activators (specific immunomodulators). Usage of the modified a-interferon as an immunomodulator was previously shown in reference [13]. Moreover, succinylated interleukin-2 has shown significantly greater activity than the native interleukin [14]. Based on these data, we hypothesized that microbial lectins - inductors of TOLL / interleukin receptors can significantly potentiate the targeted cytotoxic effect of fRNA through the activation of anticancer immunity.

Additionally to anticancer activity *in vitro* in Erlich carcinoma cells models and L1210 lymphoid leukemia cells, fRNA has shown high efficacy *in vivo* in Lewis carcinoma model of C57BL mouse lines resulting in life span of mice.

The aim of our study is the synthesis, structure confirmation and researching the anticancer activity of complementary oligo RNA, capable to block the tRNA in the cancer cell. We used the purified alanine- tRNA (atRNA) of yeasts *Saccharomyces torula* for the synthesis. atRNA was fragmented by ribonuclease, and the resulting oligonucleotides mixture was modified according to exocyclic aminogroups and ribose residues by their succinic anhydride acylation. A combination of fRNA and the inductor of TOLL-receptors based on lectin *B. subtilis* B-7025 have been proposed for usage.

## Materials and Methods

The RNA from *Saccharomyces torula* (Sigma, USA), ribonuclease (R6513, Sigma) with an activity of ≥ 70 Kunitz units / mg of protein has been used in studies. Also we used Sephadex G50- based affinity column (Pharmacia, EU) conjugated by glutar dialdehyde (Lyaoyanghengyechemical, China) by alanine amino acid (Biorad, USA) for preparative isolation of atRNA. To modify the RNA fragments we used succinic anhydride (Fluka, EU). Also for tRNA preparative extracting from affinity column and further purification, we used the following eluents: a 0.2 M phosphate, 0.5 M acetate and 0.1 M TRIS-buffers (BIORAD, USA-Israel). The product analysis was performed using ion-pair reverse-phase HPLS according to [15] with A eluent: lithium perchlorate and perchloric acid (4 M and 0.2 M, respectively) and acetonitrile as eluent B (manufactured by Econova, RF) [16]. The HPLC analysis was carried out on MilichromA-02 chromatograph in the multi-wavelength detection mode. For detecting the absence/presence of the RNA fragments at the affinity column exit we used automatic UV spectrophotometer SF-56 (Lomo, RF) at 260 nm wavelength.

### Preparation of the affinity column for a tRNA isolation [17]

The affinity column 20 mm (diameter) x 120 mm (length) was used for atRNA isolation. After swelling of 250 g Sephadex G-50 granules in 0.1 TRIS-buffer, the buffer solution was decanted. The granules were filled with a 0.3% solution of glutar dialdehyde in 40% ethanol for 15 minutes at 50°C. After this step the solution was decanted and added by 5% alanine solution in acetate buffer for 20 minutes at room temperature. Thereafter, granules in the alanine solution were charged onto the column and the unbound alanine was eluted from the column by acetate buffer of three times volume V_0_.

### *Succinylated fragmented RNA synthesis* was performed in the following manner

1.0 g of total RNA from *Saccharomyces torula* (Sigma, USA) was dissolved in 0.5M acetate buffer to a 10% concentration. Then the concentrate was passed through a previously prepared affinity column, washed by five-fold volume of pure acetate buffer. The identification of atRNA sorption completion was fixed by the absence of the signal at the exit at 260 nm. After sorbtion atRNA was eluted by phosphate buffer solution until the signal has appeared at 260 nm. Elution time was about 16 minutes. The atRNA concentration was determined using spectrophotometry by peak area at 260 nm after the sample analysis on HPLC Millichrome A-02. Previously diluted ribonuclease in phosphate buffer (RNAse/atRNA relation was 1: 100) was added to the obtained solution and incubated for 3 hours in the thermostat at 37°C. Following incubation, the mixture was again analyzed on HPLC Millichrome A-02 with determination of components concentrations.

HPLC conditions: eluent A: 4MLiClO_4_ + 0.2MHClO_4_, eluent B: CH_3_CN, gradient from 0 to 70% CH_3_CN, column thermostat temperature 60°C, and multi-wavelength detection. Succinylation of the atRNA oligomeric fragments was performed as shown in reference. fRNA finished product has undergone HPLC similarly as for the unmodified fragmented atRNA. The resulting solution was filled into vials and sealed. These vials were then sterilized by autoclaving for 15 min at 120°C. The average RNA concentration in the solution by the peak areas sum was 8.0 ± 0.2%. The obtained solution was further used to examine the anticancer activity *in vitro* and *in vivo*. For the immunotherapy we used immunotropic 70 kDa lectin *B. subtilis B-7025* (Vaccine) (0.3 mg / ml). [18,19]

Experiments were carried out on C57Bl line mice (males, 2.5 months, 20-22g from vivarium at Kavetsky IEPOR, National Academy of Sciences of Ukraine, Kyiv, Ukraine). Animal care and handling was carried out in accordance with generally accepted international rules of research in experimental animals. Lewis lung carcinoma (LLC) was used as an experimental model. The LLC was inoculated intramuscularly in the hind limb thigh (3x10^5^ cells per animal).

The vaccination started on the sixteenth day after tumor inoculation. The vaccine was administered subcutaneously in the animals’ withers (16, 19, 26 and 33 day of tumor growth) at the rate of 0.3 ml per injection. The start of fRNA administration began on day 20 after tumor inoculation, intraperitoneally for five consecutive days, and continued to sevenfold administering once every three days, at the rate of 5 μl of the diluted solution per animal (0.3 ml of physiological sodium chloride solution). The control group received intraperitoneal cisplatin injections of 50 μl for five consecutive days, starting from the day 20. The following groups have been formed: 1 control of tumor growth (CTG) - animals with inoculated LLC tumor; 2 cisplatin - animals which received cisplatin; 3 fRNA - animals which received fRNA; 4 Vaccine - animals which received KPV; 5 fRNA + vaccine - animals that received fRNA and vaccine. During the experiment, the dynamics of tumor growth was assessed.

Antimetastatic effect was evaluated on the fortieth day after LLC inoculation based on number and size of lung metastases: incidence of metastasis (%); the average number of metastases in a mouse; the average volume of metastases per a mouse. The cytotoxic activity (CTA), lymphocytes (L), and macrophages (Mf) were determined *in vitro* using the MTT assay [20]. The LLC cells were used as target cells.

The statistical analysis of the results was performed by means of commonly accepted variation statistics methods using analysis of variance. The study results are presented as M ± m, where M is the arithmetic mean, m is its standard error. Differences were evaluated as valid if p <0,05. The calculations and graphing were performed using MS Excel application.

## Results and Discussion

In our first studies of acylated polynucleotides, a new phenomenon was identified: polynucleotide,s acylated on exocyclic amino groups, selectively hybridized only with their nonacylated predecessors [21]. Insoluble, infusible conjugates were formed under such conditions. That was exactly infusible feature that differed classic double-stranded DNA from the obtained hybrids.

They did not melt at any temperature, and did not dissolve in anything other than concentrated alkalis. We explain this feature of the synthesized acyl-DNA as the change of the bond forming principle between the DNA chains: from the hydrogen to the hybrid chain: hydrogen-ionic bond. This chemical bond is absolutely not provided by natural DNA repair mechanisms. Thus, nuclease and cellular helicases are ineffective in forming the hybrids between such acyl-DNA (RNA) and their non-acylated precursors.

Additionally to polynucleotides, we have shown dependency of charge/activity in some other acylated biopolymers: proteins, polysaccharides, polynucleotides, tannins, bacteriophages and immunoglobulins. Based on these studies, we have developed and implemented Albuvir, a new veterinary antiviral medicinal product [22]. We have found that a certain precision modification of biopolymer structure is capable to increase its biological activity, or completely change its features, or result in formation of self-assembled structures. Detected patterns served as the basis of the principle of obtaining microRNA with acylated exocyclic amino groups.

We used the ensemble of oligonucleotides, which were obtained by polynucleotides hydrolysis, but with reverse changes of molecular charges. The ensemble is a term from supramolecular chemistry. The objects of supramolecular chemistry are supramolecular ensembles, spontaneously built from complementary fragments, that have geometric and chemical matching fragments, like the spontaneous assembly of complex spatial structures in living cells [23].

This assembly for fRNA occurs by attaching of fRNA fragments to the whole aRNA, while there is a selective irreversible conjugation between 8 fragments of fRNA and the whole aRNA in a cancer cell with a complex denaturation and agglutination, like a similar reaction between pUC18 and pBR322 plasmids with succinylated derivatives [24]. Fig. 1 demonstrates a chromatogram of oligonucleotides mixtures after aRNA ribonuclease processing.

**Fig.1.**
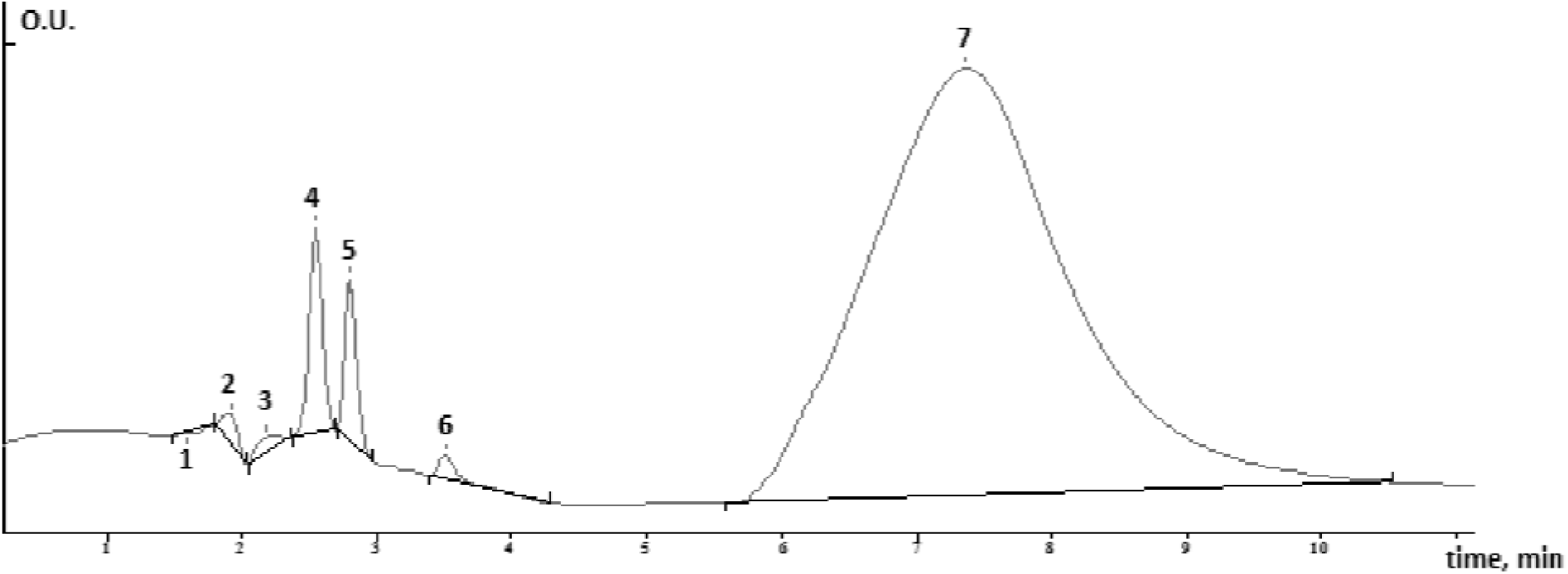
Chromatogram of the oligonucleotides mixture disintegration – products of aRNA ribonuclease processing; chromatographic conditions: eluent A: 4MLiClO4 + 0.2 MHClO_4_, eluent B: CH_3_CN, gradient from 0 to 70% CH_3_CN, column thermostat temperature of 60°C, multi-wavelength detection.

According to Fig. 1, the baseline content has seven components corresponding to UV spectra of the RNA, where peak No. 7dominates with the maximum retention time from 6 to 9 minutes. Fig. 2 shows a chromatogram of the final product division after the chemical modification.

**Fig.2.**
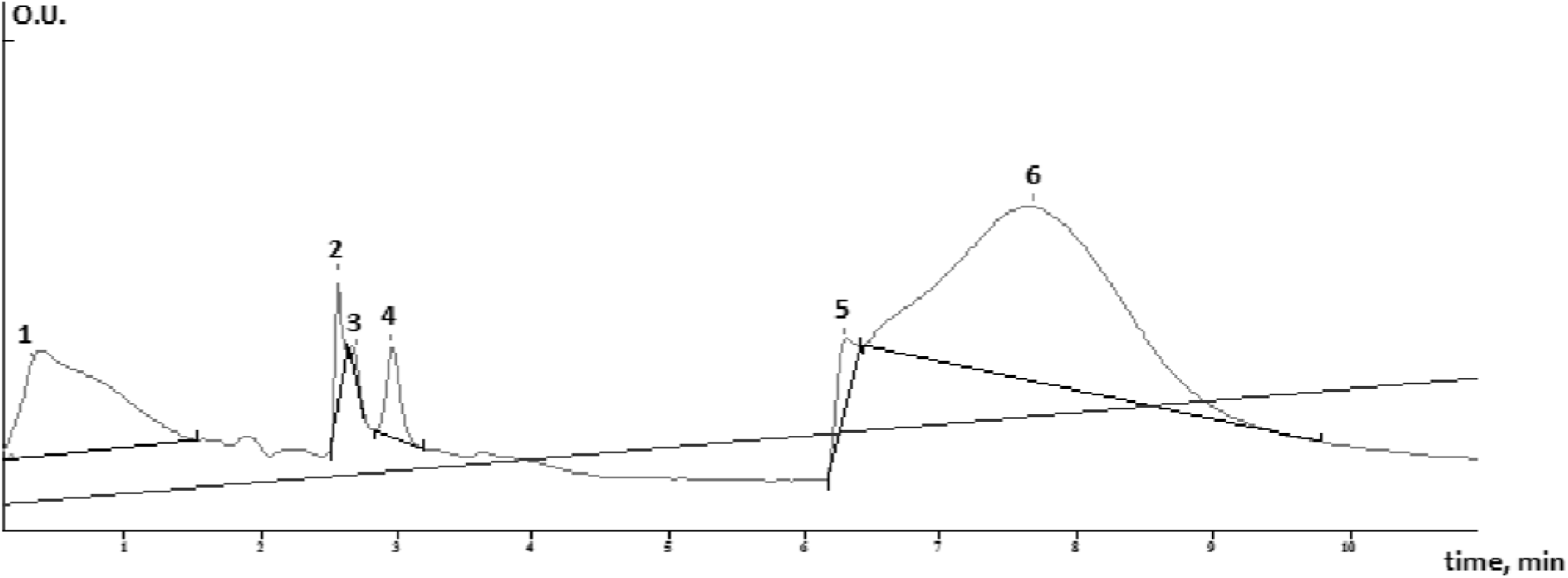
Chromatogram the fRNA (final product); chromatographic conditions: eluent A: 4MLiClO_4_ + 0.2 MHClO_4_, eluent B: CH_3_CN, gradient from 0 to 70% CH_3_CN, column thermostat temperature 60°C, multi-wavelength detection.

The fRNA (final product) contained 3 additional peaks No. 1, 2 and 5 on the chromatogram (Fig. 2).

UV spectra of all peaks of the primary hydrolyzate (Fig. 1.) were identical and consistent with the integral UV spectrum of A + G + C + V (Fig. 3).

**Fig. 3.**
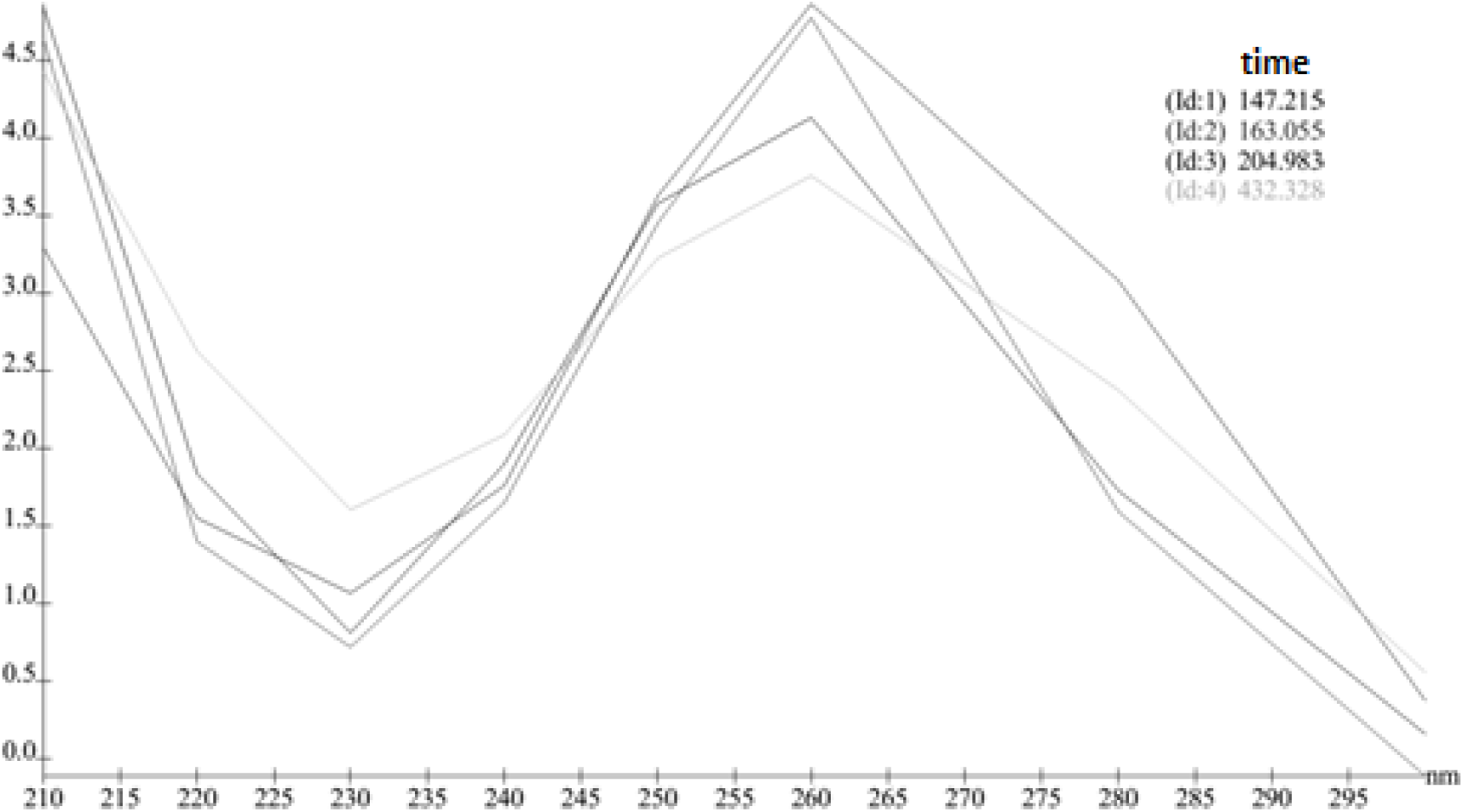
The integrated electronic absorption spectrum in the UV range from 210 to 300 nm for all peaks for the of the baseline atRNA-hydrolyzate chromatogram: nearly all peaks spectra coincide, which confirms that this is exactly RNA fragments, but they have different molecular weight.

Fig. 4 shows integrated UV–spectra for the final product – succinylated hydrolyzate of atRNA (fRNA).

**Fig. 4.**
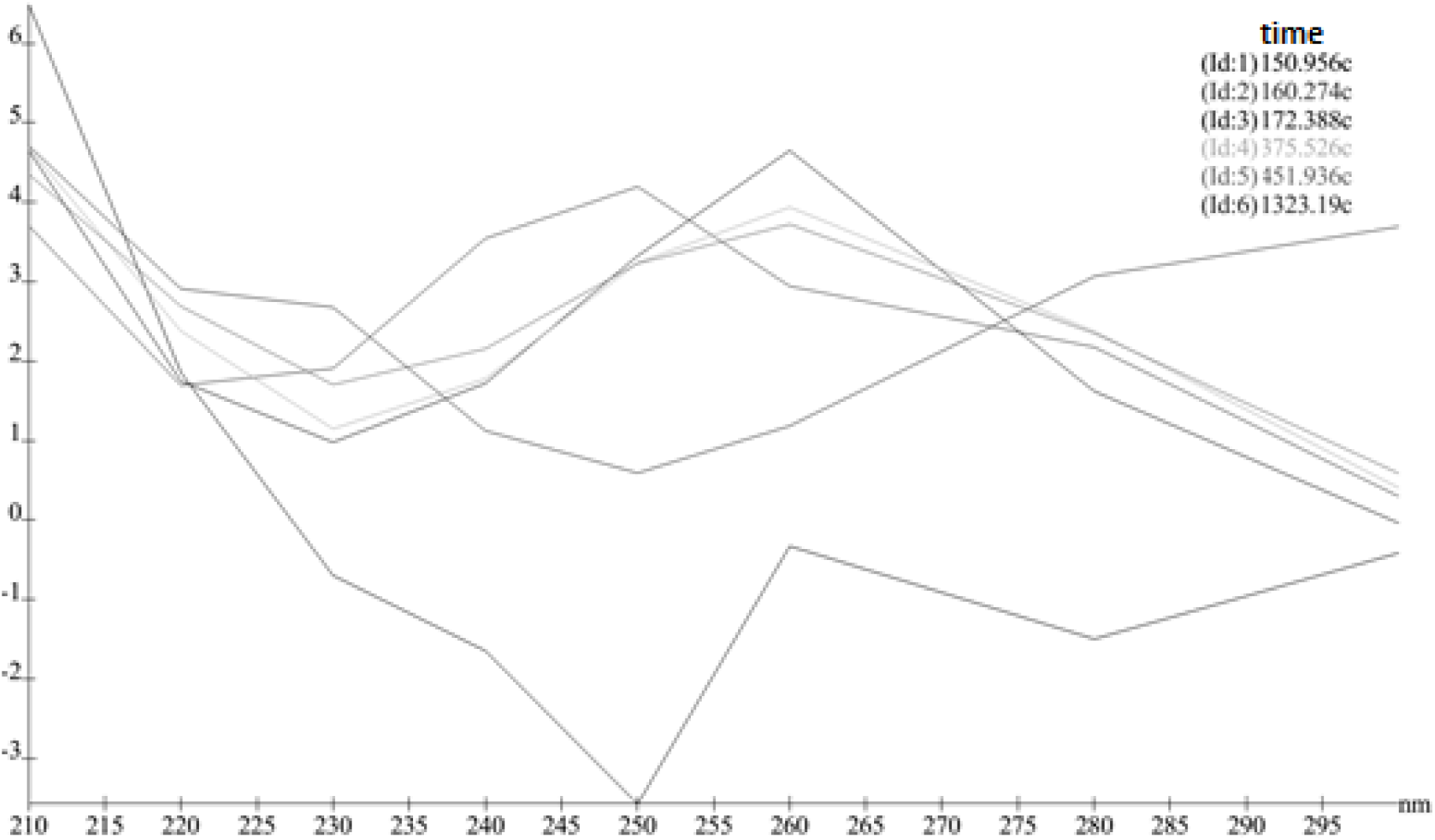
The integrated electronic absorption spectrum in the UV range from 210 to 300 nm from the chromatogram all fRNA peaks: the spectra of new peaks significantly differ from the spectra of the primary atRNA-hydrolyzate, which confirms completion of the reaction of RNA oligonucleotides structure modification.

According to Fig 4 the spectral correlation of the majority of peaks has been significantly changed. This effect supports the hypothesis about significant change in the structure of the finished product relative to the primary one. In addition, there have appeared three additional asymmetric peak in the chromatogram with a retention time of 0.3; 2.6; 6.4 min.

On average, alanine tRNAs from the all RNA mass is about 1% out of the total amount of RNA. There are significant differences between chromatograms of fRNA and the baseline fragmented (Fig. 4) (but not modified) RNA, which allow standardizing of the finished product and developing on its basis, the new pharmaceutical formulations and dosage forms.

The study of fRNA antitumor activity in its complex usage with a microbial lectin (Vaccine) in mice with Lewis lung carcinoma (LLC).

When evaluating the animals’ survival, it was found that at day 41 (end point of the experiment) in the group of animals, which received the fRNA combined therapy with the antitumor vaccine, all study animals were alive, whereas in the control group at the time of the study only 15% of animals were live. Animal survival rate in groups receiving either the fRNA or Vaccine was 70 and 40%, respectively. In the Cisplatin group of animals the survival rate was 30%.

**Fig.5.**
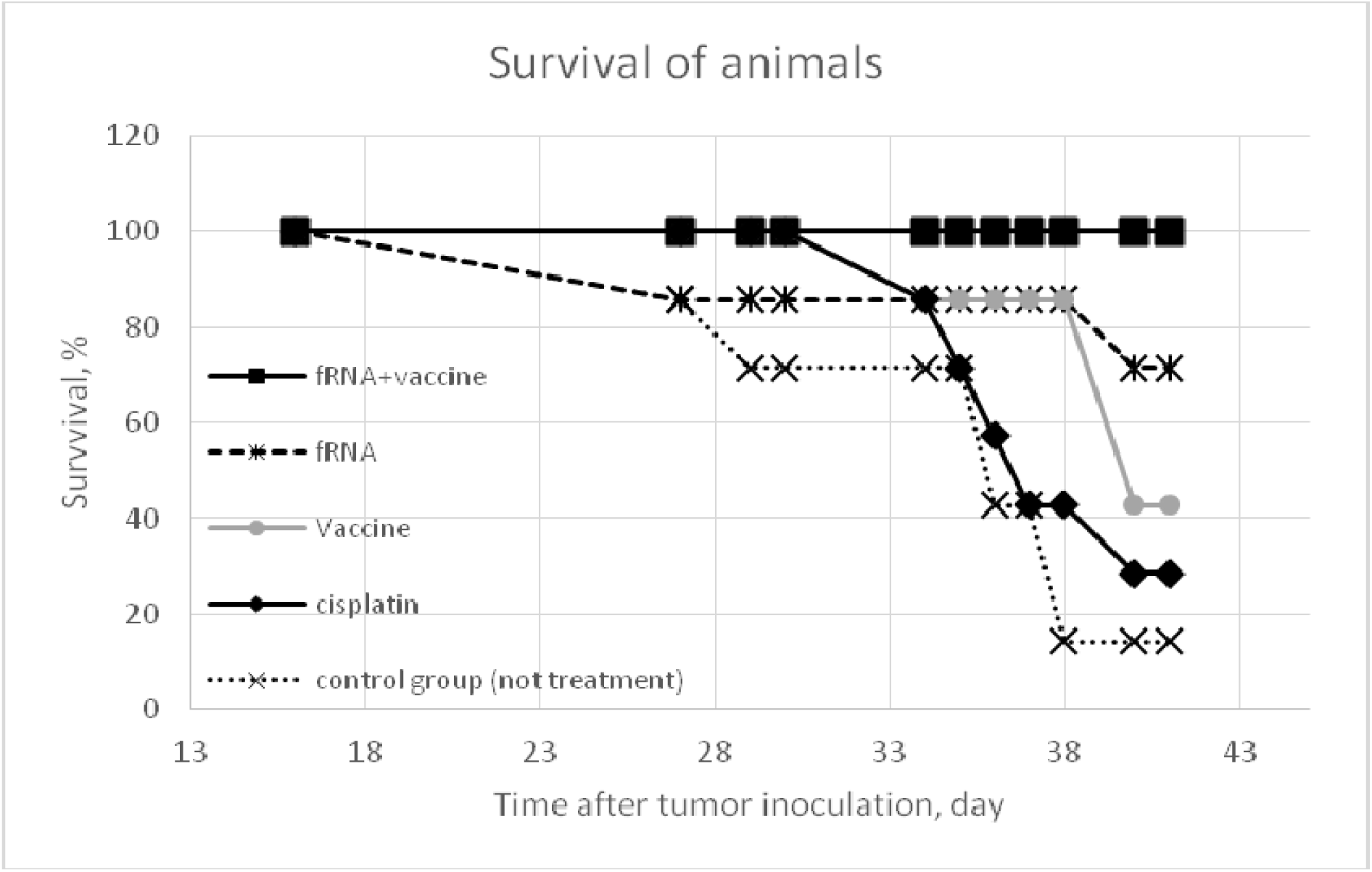
Survival of animals

When studying the tumor growth dynamics, it was found that in the animals group treated with the combined therapy fRNA and Vaccine, the tumor size was significantly (p ≤0,05) lower compared to the control during the entire study period. The animals group, treated with fRNA in monotherapy, and also during the study period, showed a primary tumor growth delay, but it was less pronounced. The efficacy of the vaccine therapy regarding the inhibition of the primary tumor node growth, coincided with the efficiency of cisplatin. By the thirty-sixth day, the average tumor volume in groups made up 7.05 ± 1.10 mm^3^- in the control group of tumor growth; 6.30 ± 2.42 **-** in “Cisplatin” group; 5.31 ± 0.76 - in the “fRNA” group; 5.50 ± 0.56 - in the “Vaccine” group and 2.61 ± 0.23 – in the “fRNA + Vaccine” group.

**Fig. 6.**
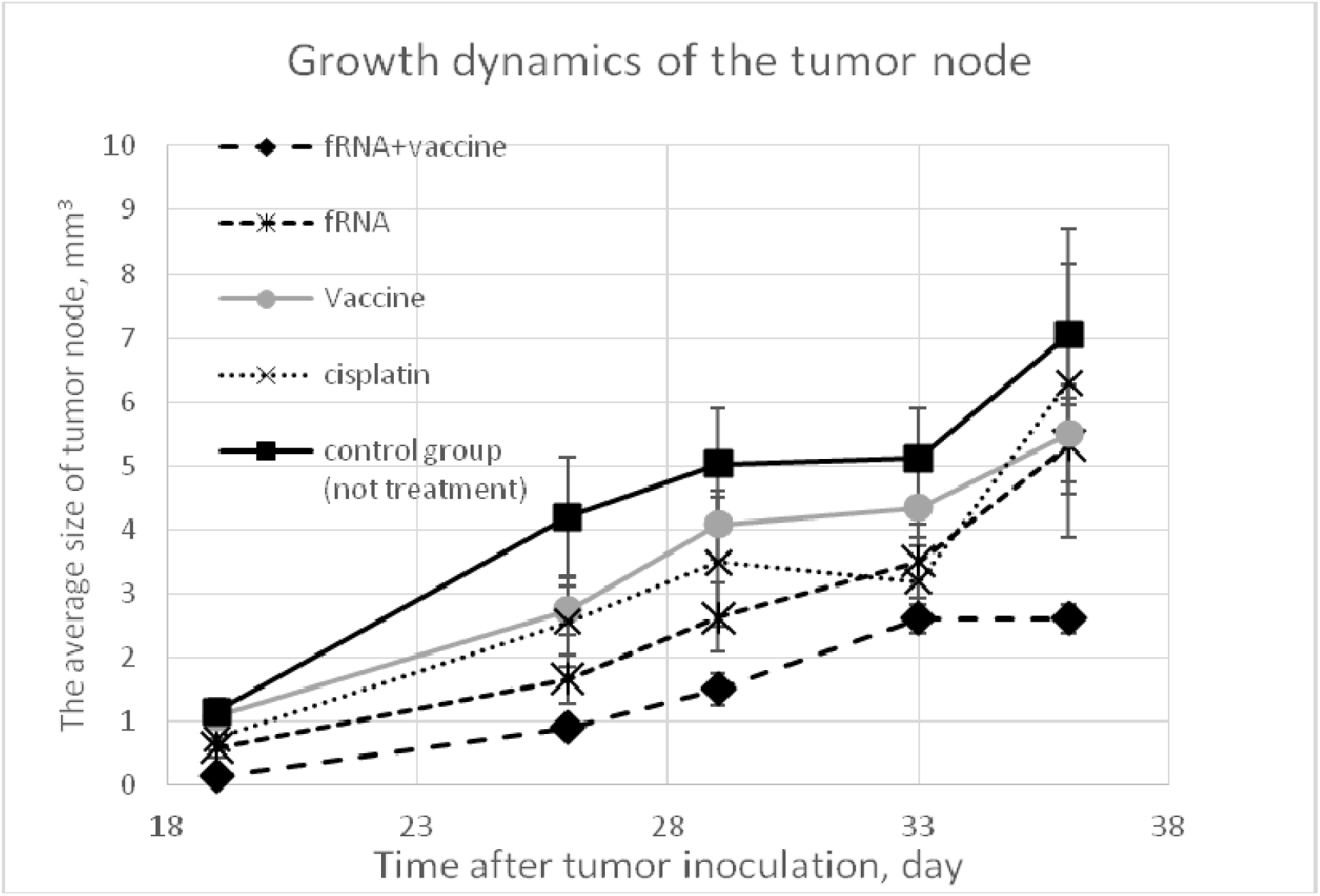
The dynamics of the tumor node growth

The therapy antimetastatic effects study demonstrated that the use of the Vaccine, fRNA, fRNA + Vaccine complex led to the development and propagation delay of metastatic nodes in the experimental animals (Table 1).

**Table 1.** Level of metastasis in experimental animals

|  | Volume mm<sup>3</sup>/animal |  | Quantity, pcs/animal |  |
| --- | --- | --- | --- | --- |
|  | M | m | M | m |
| Control | 339.958 | 0 | 159 | 0 |
| fRNA | 70.99879 | 19.94599 | 46.14286 | 9.43254 |
| Vaccine | 55.7245 | 24.90542 | 35 | 13.778 |
| Vaccine+<br>fRNA | 60.2463 | 25.1966 | 45.4 | 17.96831 |
| Cisplatin | 10.21875 | 3.90775 | 13 | 3 |

When evaluated by the average life expectancy (Table 2), it was found that in the group of animals, treated with the combined therapy aRNA/Vaccine, was the largest and lasted for 41 days, whereas the use of both types of therapy in mono mode reduced the average life span in animals up to 38 days.

**Table 2.** The average life expectancy (ALE) in the experimental animals

|  | ALE |  |
| --- | --- | --- |
|  | M | m |
| Control | 34.42857 | 1.83689 |
| Cisplatin | 36.42857 | 1.49375 |
| fRNA/Vaccine | 41 | 0 |
| Vaccine | 38.14286 | 1.4708 |
| fRNA | 38.57143 | 1.97432 |
| Intact mouse | -- | -- |

When evaluated the immunological effects of the therapy (Table 3.), we found that use of either fRNA or vaccines resulted to the increase of the direct cytotoxic lymphocyte activity compared with other groups. The cytotoxic activity of macrophages in all treated animals was at the level of intact animals (lower than activity indicators in the control group of tumor growth). In the group of animals treated only with Vaccine the direct cooperative cytotoxicity of lymphocytes and macrophages was slightly increased.

**Table 3.** Indicators of immune cell cytotoxic activity in the MTT test (cytotoxicity index %)

| MTT | Control,<br>M±m | Cisplatin<br>M±m | Vaccine/<br>fRNA<br>M±m | Vaccine<br>M±m | fRNA<br>M±m | Intact<br>mouse<br>M±m |
| --- | --- | --- | --- | --- | --- | --- |
| CTL* | 40.68±1.86 | 49.51±1.70 | 46.14±4.46 | 63.61±2.29 | 67.77±1.08 | 37.81±0.09 |
| CTM** | 48.14±0.93 | 28.77±1.41 | 37.55±0.31 | 31.41±4.20 | 26.21±2.01 | 32.46±5.70 |
| CCT***<br>(L+M) | 65.75±2.58 | 68.91±2.79 | 65.15±2.03 | 84.40±2.47 | 57.13±0.77 | 67.98±2.63 |

\* - cytotoxicity of lymphocytes; \*\* - cytotoxicity of macrophages; \*\*\* - cooperative cytotoxicity

Combined use of fRNA and Vaccine has the advantage compared with the use of these drugs in monotherapy, as the anticancer efficacy of the scheme is much higher, which is manifested in the primary tumor node and metastasis inhibition.

## Conclusion

The fRNA (final product) contained 3 additional peaks No 1, 2 and 5 on the chromatogram. UV spectra of all peaks of the primary atRNA- hydrolysate were identical and consistent with the integral UV spectrum of A + G + C + V. When evaluating the animals’ survival, it was found that at day 41 (end point of the experiment) in the group of animals, which received the fRNA combined therapy with Vaccine, all study animals were alive, whereas in the control group at the time of the study only 15% of animals survived. Animal survival rate in groups receiving either the fRNA or Vaccine was 70 and 40%, respectively. In the Cisplatin group of animals the survival rate was 30%. In the therapy antimetastatic effects study it was found that the use of Vaccine, fRNA, fRNA + Vaccine complex resulted to the development and propagation delay of metastatic nodes in the experimental animals. Combined use fRNA and Vaccine has the advantage compared with the use these drugs in monotherapy, as the anticancer efficacy of the scheme is much higher, which is manifested in the primary tumor node and metastasis inhibition.

## Acknowledgements

This work was supported in part by the Farber Center of Academic Success (New-York, USA), clinic “Modern Medical Group” (New-York, USA), the National Academy of Sciences of Ukraine, the National Academy of Medical Sciences of Ukraine.

## References

[1] Cross, D.; Burmester, J.K. Gene Therapy for Cancer Treatment: Past, Present and Future. Clin. Med. Res., 2006, 4(3), 218–227,

[2] Rayburn, E.R.; Zhang, R. Antisense, RNAi, and Gene Silencing Strategies for Therapy: Mission Possible or Impossible? Drug. Discov. Today., 2008, 13(11-12), 513–521.

[3] Mansoor, M.; Melendez, A.J. Advances in Antisense Oligonucleotide Development for Target Identification, Validation, and as Novel Therapeutics. Gene Regul. Syst. Bio., 2008, 2, 275–295,

[4] Juliano, R.; Alam, M.R.; Dixit, V.; Kang, H. Mechanisms and strategies for effective delivery of antisense and siRNA oligonucleotides. Nucleic Acids Res., 2008, 36(12), 4158–4171.

[5] Falzarano, M.S.; Passarelli, C.; Ferlini, A. Nanoparticle Delivery of Antisense Oligonucleotides and Their Application in the Exon Skipping Strategy for Duchenne Muscular Dystrophy. Nucleic Acid Ther., 2014, 24(1), 87–100

[6] Moreira, J.N.; Santos, A.; Moura, V.; Pedroso de Lima, M.C.; Simoes, S. Non-viral lipid-based nanoparticles for targeted cancer systemic gene silencing. J Nanosci Nanotechnol., 2008, 8(5):187-2204.

[7] Lu, J.J.; Langer, R.; Chen, J. A novel mechanism is involved in cationic lipid-mediated functional siRNA delivery. Mol. Pharm. 2009, 6(3),763-771.

[8] Allen, T.M. Ligand-targeted therapeutics in anticancertherapy. Nat. Rev. Cancerv.,2002, 2, 750–763.

[9] Bao, G.; Bao, X.R. Shedding light on the dynamics of endocytosis and viral budding. Proc Natl Acad Sci USA., 2005, 102, 9997–9998.

[10] Gabizon, A.; Shmeeda, H.; Barenholz, Y. Pharmacokineticsof pegylated liposomal doxorubicin – review of animaland human studies. Clin. Pharmacokinetics.,2003, 42, 419–436,

[11] Gao, H.; Shi, W.; Freund, L.B. Mechanics of receptor-mediated endocytosis. Proc Natl Acad Sci USA., 2005, 102(27), 9469-9474.

[12] Martynov, A.V.; Bomko, T.V.; Nosalskaya, T.N.; Farber, B.S.; Farber, S.B. Oral long-acting pharmaceutical form of insulin on the basis of self-organizing kvasi-living system of combinatorial peptides. Annals of Mechnikov Institute., 2012, 2, 64-70.

[13] Martynov, A.V.; Smelyanskaya, M.V. Antiproliferative properties of chemically modified - 2b-recombinant interferon. J of Interf. & Cytokine res.,2005, 25 (7): 414-417.

[14] Martynov, A.V.; Farber, B.S.; Farber, S.B.; Kabluchko, T.V. Synthesis of the ensembles from succinylated interleukin-2 derivatives and their biological activity in vitro. ScienceRise., 2015, 11 (4): 25-30.

[15] Gilar, M.; Fountain, K.J.; Budman, Y; Neue, U.D.; Yardley, K.R.; Rainville, P.D.; Russell, II R. J.; Gebler J.C. Ion-pair reversed-phase highperformance liquid chromatography analysis of oligonucleotides: Retention prediction. J. Chromatogr., 2002, 958 (1-2), 167-182.

[16] Baram, G.I.; Grachev, A. Use of lithium perchlorate for isolation and analysis of oligo- and polynucleotides. Bioorg chem., 1985, 11(10), 1420-1422.

[17] Martins, Ana Rita Nunes. Biorecognition by amino acid-based affinity chromatography for RNA purification. Diss. UNIVERSIDADE DA BEIRA INTERIOR, 2013.

[18] Didenko, G.V.; Sorokina, L.V.; Shpak, Eu. G. The application of fullerene C60 for the modification of an anticancer vaccine based on metabolism products of Bacillus subtilis 7025. Journal of Biological Physics and Chemistry., 2011, 11, 30–35.

[19] Potebnya, G.P.; Tanasienko, O.A.; Titiva, G.P. Specificity and biological activity of cytotoxic lectins synthesized by Bacillus subtilis B-7025. Exp Oncol., 2002, 2, 250–252.

[20] Ohno, M.; Abe, T. Rapid colorimetric assay for the quantification of leukemia inhibitory factor (LIF) and interleukin-6 (IL-6). J Immunol Meth., 1991, 145, 199–203.

[21] Martynov, A.V. Synthesis and properties of succinylated polynucleotides. News in Pharmacy (Ukr)., 2003, 2(32), 20-23.

[22] Martynov, A.V.; Babkin N.; Zheynova N. Antiviral activity of the albuvir in models vesicular stomatitis virus and human herpes virus type 1 in vitro. Scientific Bulletin of Lviv National University of Veterinary Medicine and Biotechnology named. Gzhytsky., 2011, 2 (1), 181-184.

[23] Jean-Marie Lehn. Supramolecular Chemistry. Concepts and Perspectives. Weinheim; New York; Basel; Cambridge; Tokyo: VCH Verlagsgesellschaft mbH., 1995, Chapter 7, 103.

[24] Patent WO2012070965A1. Modified antisense oligopeptides with anticancer properties and method of obtaining them. Martynov A, Farber B, Farber S. /Appl. US 12/931,468; 25 Mat 2012, Priority 22 Nov 2010

